# Differential regulation of gene expression pathways with dexamethasone and ACTH after early life seizures

**DOI:** 10.1101/2022.04.21.489067

**Authors:** Jeffrey L. Brabec, Mohamed Ouardouz, J. Matthew Mahoney, Rod C. Scott, Amanda E. Hernan

**Affiliations:** University of Vermont, Department of Neurological Sciences, 149 Beaumont Avenue, Burlington, VT 05401; Nemours Children’s Health, Division of Neuroscience, 1600 Rockland Road, Wilmington, DE 19803; The Jackson Laboratory, 600 Main Street, Bar Harbor, ME 04609; Neurosciences Unit University College London, Institute of Child Health, London WC1N 1EH, UK; University of Delaware, Psychological and Brain Sciences, South College Avenue, Newark, DE 19716, USA

**Keywords:** ACTH, dexamethasone, ELS, RNAseq

## Abstract

Early-life seizures (ELS) are associated with persistent cognitive deficits such as ADHD and memory impairment. These co-morbidities have a dramatic negative impact on the quality of life of patients. Therapies that improve cognitive outcomes have enormous potential to improve patients’ quality of life. Our previous work in a rat flurothyl-induction model showed that administration of adrenocorticotropic hormone (ACTH) at time of seizure induction led to improved learning and memory in the animals despite no effect on seizure latency or duration. Administration of dexamethasone (Dex), a corticosteroid, did not have the same positive effect on learning and memory and has even been shown to exacerbate injury in a rat model of temporal lobe epilepsy. We hypothesized that ACTH exerted positive effects on cognitive outcomes through beneficial changes to gene expression and proposed that administration of ACTH at seizure induction would return gene-expression in the brain towards the normal pattern of expression in the Control animals whereas Dex would not. Twenty-six Sprague-Dawley rats were randomized into vehicle- Control, and ACTH-, Dex-, and vehicle-ELS. Rat pups were subjected to 60 flurothyl seizures from P5 to P15. After seizure induction, brains were removed and the hippocampus and PFC were dissected, RNA was extracted and sequenced, and differential expression analysis was performed using generalized estimating equations. Differential expression analysis showed that ACTH pushes gene expression in the brain back to a more normal state of expression through enrichment of pathways involved in supporting homeostatic balance and down-regulating pathways that might contribute to excitotoxic cell-damage post-ELS.

## Introduction

Seizures that occur during early life development are associated with persistent cognitive abnormalities including depression, ADHD, autism spectrum disorders and memory impairments (Dunn and Kronenberger, 2005; Kanner, 2005; Matson et al., 2010; Rantanen et al., 2010). These co-morbid symptoms frequently have a negative impact on quality of life and therefore deserve treatment. To date, treatment approaches have largely targeted seizure outcomes with the view that preventing seizures will ameliorate the cognitive deficits that accompany them (Nariai et al., 2018). In clinical practice, however, prevention of seizures has an unreliable impact on cognitive co-morbidities (Nariai et al., 2018). Therapies that alter cognitive outcomes, even in the absence of altering seizure outcomes, therefore may have enormous potential for improving quality of life.

Immature rats that are exposed to 50-60 flurothyl-induced generalized seizures have long-term cognitive impairments. We have previously made the remarkable observation that administration of ACTH at the time of seizure induction has no effect on latency to seizure or duration of seizures, but resulted in improved learning and memory (Massey et al., 2016). ACTH is a drug given to patients with severe epilepsy that is often grouped with other corticosteroids and presumed to exert its actions through targeting corticoid receptors systematically to suppress inflammation. However, in our previous work, the administration of dexamethasone (Dex), a corticosteroid, did not have a similar effect at preventing cognitive impairment in our rat model (Massey et al., 2016) and was even shown to exacerbate injury in a rat model of temporal lobe epilepsy (Duffy et al., 2014). This suggests that ACTH is exerting its positive effect via pathways that are unrelated to seizure prevention and are unlikely to be the result of only direct endocrine effects such as that exerted by corticosteroids.

The full complement of diverse epilepsy symptoms arises from insults to the underlying functional and genetic networks. After a seizure, pro-inflammatory factors, pro-senescence messengers, and glial activation genes all show an increase in expression from baseline, while pro-survival pathways are down-regulated in damaged neurons (Ghosh et al., 2022). This altered expression state across the brain underlies dysregulation of the neural network that generates seizures and drives cognitive dysfunction. An ideal treatment would target these gene expression trajectories by modulating pathways to prune irreparably damaged and dying neurons and promote survival pathways in those still remaining in order to return the brain to a normal gene expression state.

Dex and ACTH differentially influence long-term cognitive outcomes, implicating distinct effects on the molecular physiology of the brain. We hypothesized that the difference in the two mechanisms would be observable in differential gene expression profiles of neural tissue from animals treated with each drug. Further, we hypothesized that animals that experienced recurrent early life seizures (ELS) treated with ACTH would have a gene expression pattern which was more similar to that of the control animals (i.e. fewer differentially expressed genes compared to control) than ELS animals treated with Dex. We then performed a detailed pathway analysis of the ACTH and control expression profiles to identify potentially targetable pathways that could lead to development of improved treatment options for cognitive deficits in patients with epilepsy.

## Materials and Methods

### Animal model

A total of 26 Sprague-Dawley pups were randomized into four groups; Controls that were separated from the dam at the time of seizures to control for maternal separation anxiety and injected with the vehicle solution, but experiencing no seizures (n=6) and 3 groups with early life seizures. The treatments were placebo (5% gelatin used to dilute ACTH; n=3), ACTH (dose 150 IU/m2; n=8), and Dex (dose 0.5mg/kg; n=3). One hour before the first seizure on each day the animals received a subcutaneous injection of drugs as above. The pups were then subjected to 60 flurothyl induced seizures (6 daily) from P5 to P15. The pups were removed from the dams and placed in a sealed container with 4 chambers. Liquid flurothyl (0.1mL), an inhaled convulsive agent, was slowly dripped onto a small piece of filter paper within the container. Pups were observed carefully, and removed from the chamber and allowed to ventilate once they showed signs of tonic forelimb and hindlimb extension. Littermate control pups were removed from the dam and handled at this time to control for the effect of handling and of_maternal separation-related stress. Seizures occurred in all animals irrespective of treatment. Two days after the last ELS induction the pups were anesthetized with isoflurane then decapitated and the brains removed and dissected in cold RNA-later (4° C). Both PFC and hippocampus were removed, and flash frozen in a dry ice ethanol bath and stored in an Eppendorf tube. All the samples were kept at −80° C until processed for RNA extraction.

### RNAseq analysis

The samples were transferred to the Vermont Integrated Genomics Core for RNA sequencing. Sample libraries were constructed using the SMARTer Stranded Total RNA-Seq kit v2 Pico Input Mammalian protocol (Takara Bio, San Jose, CA). Paired-end sequencing was performed using an Illumina HiSeq system (Bentley et al., 2008). Before downstream analysis, sequences were demultiplexed and had adapters removed.

### Gene expression modeling

After post-sequencing quality control, samples were transferred to and processed on the Vermont Advanced Computing Cluster (VACC). Sample read files were assessed for quality using FastQC and reads were trimmed using cutadapt (Martin, 2011). Next, reads were aligned to the latest release of the rat genome from Ensembl (version 102) (Yates et al., 2019) using the STAR alignment tool (Dobin et al., 2013). Aligned sequences then underwent quality control analysis using Picard Tools (http://broadinstitute.github.io/picard/) to determine alignment quality and to identify any duplicate sequences that may have been present. All samples that passed QC and gene counts were quantified using HTSeq (Anders et al., 2015). Count data for each gene was merged into a counts table for the whole dataset and imported into the statistical programming language, R, for differential expression analysis.

Differential expression pre-processing was conducted using the DESeq2 R package (Love et al., 2014). The counts matrix was read into R along with a metadata table containing information on the experimental groups (Control, Dex, ACTH, ELS) and sex of the animals, as well as the brain locations from which the samples were obtained. Genes with fewer than ten counts across all samples were filtered out and were excluded from further analysis. The data was normalized and transformed using a regularized log function.

The differential expression analysis was performed using generalized estimating equations (GEEs) with sex of the animal and brain region from which each sample was extracted as covariates, as well as an interaction term of treatment group by brain region. This analysis was run twice, once with ‘Control’ as the baseline comparator and once with ‘vehicle ELS’ as the baseline comparator. This allowed us to make comparisons between each of our drug treatment groups to both groups. P-values were corrected using False-Discovery Rate (FDR) and significant genes were identified as those having a corrected p-value < 0.1.

### Pathway analysis

Pathway analysis was conducted using the gost function from the gprofiler2 R package (Kolberg et al., 2020). DEGs from each model contrast were submitted to the gost function for analysis which uses a hypergeometric test to identify significantly enriched GO terms. Pathways were considered significantly enriched if they reached an FDR corrected p < 0.05. Semantic plots were generated by first using REVIGO to consolidate semantically similar GO terms into a representative term then plotting the semantic space in R (Supek et al., 2011).

### Experimental design and Statistical Analyses

Twenty-six Sprague-Dawley rats were used in this study (n = 13 males, n = 10 females). These animals were split into four groups: Control (n = 5 males, n = 4 females), ACTH (n = 6 males, n = 3 females), Dex (n = 3 females), ELS (n = 2 males, n = 3 females). Brain punches were taken from the hippocampus and prefrontal cortex of each animal RNAseq counts data was normalized using a regularized log function and differential expression was calculated using Generalized Estimating Equations. P-values were corrected using a false-discovery rate (FDR) correction. Enriched pathways were identified using a hypergeometric test with an FDR correction.

### Code Accessibility

All code to reproduce the analysis of this project can be found in the GitHub repository for this paper at https://github.com/MahoneyLabGroup/acth_deg.

## Results

Each animal had 60 seizures over the course of 10 days. We have previously reported no changes in seizure intensity or duration with treatment, but improvement in cognitive outcome (Massey et al., 2016). We chose to examine differential expression in the PFC and the hippocampus because of their relationship to aspects of cognition that are altered after ELS.

### Gene Expression in ACTH-treated rats aligns with that of Control rats

We performed differential gene expression (DEG) analysis using two GEE models. In the first model, the ‘Control’ group was set as the baseline comparator and resulted in three contrasts that we explore below: ACTH vs. Control, Dex vs. Control, and vehicle ELS vs. Control. In the second model, the ‘vehicle ELS’ group was set as the baseline comparator resulting in three additional contrasts: ACTH vs. vehicle ELS, Dex vs. vehicle ELS, and Control vs. vehicle ELS. This two-model approach allowed us to evaluate whether Dex or ACTH would normalize differential expression after ELS by looking at expression differences compared to the “normal” condition and the untreated ELS condition.

We hypothesized that animals treated with ACTH would have fewer DEGs compared to baseline Control animals than those treated with Dex. We further hypothesized that the ACTH vs. vehicle ELS contrast would share a larger proportion of DEGs with the Control vs. vehicle ELS contrast than the Dex vs. vehicle ELS contrast. The Venn diagrams in Figure 1 show that animals treated with ACTH had a total of 1,504 DEGs in the baseline Control model, and those treated with Dex had 1,693 DEGs, indicating a somewhat dissimilar pattern of gene expression to the animals without seizures, which approached significance (Chi-squared *p* = 0.054). Despite a large proportion of genes in the ACTH context with differential expression compared to Control, the two cohorts shared 517 DEGs when compared to ELS while Dex only shared 344 DEGs with Control, despite having a greater number of total DEGs compared to ELS (Chi-squared *p* = 10e-13).

**Figure 1:**
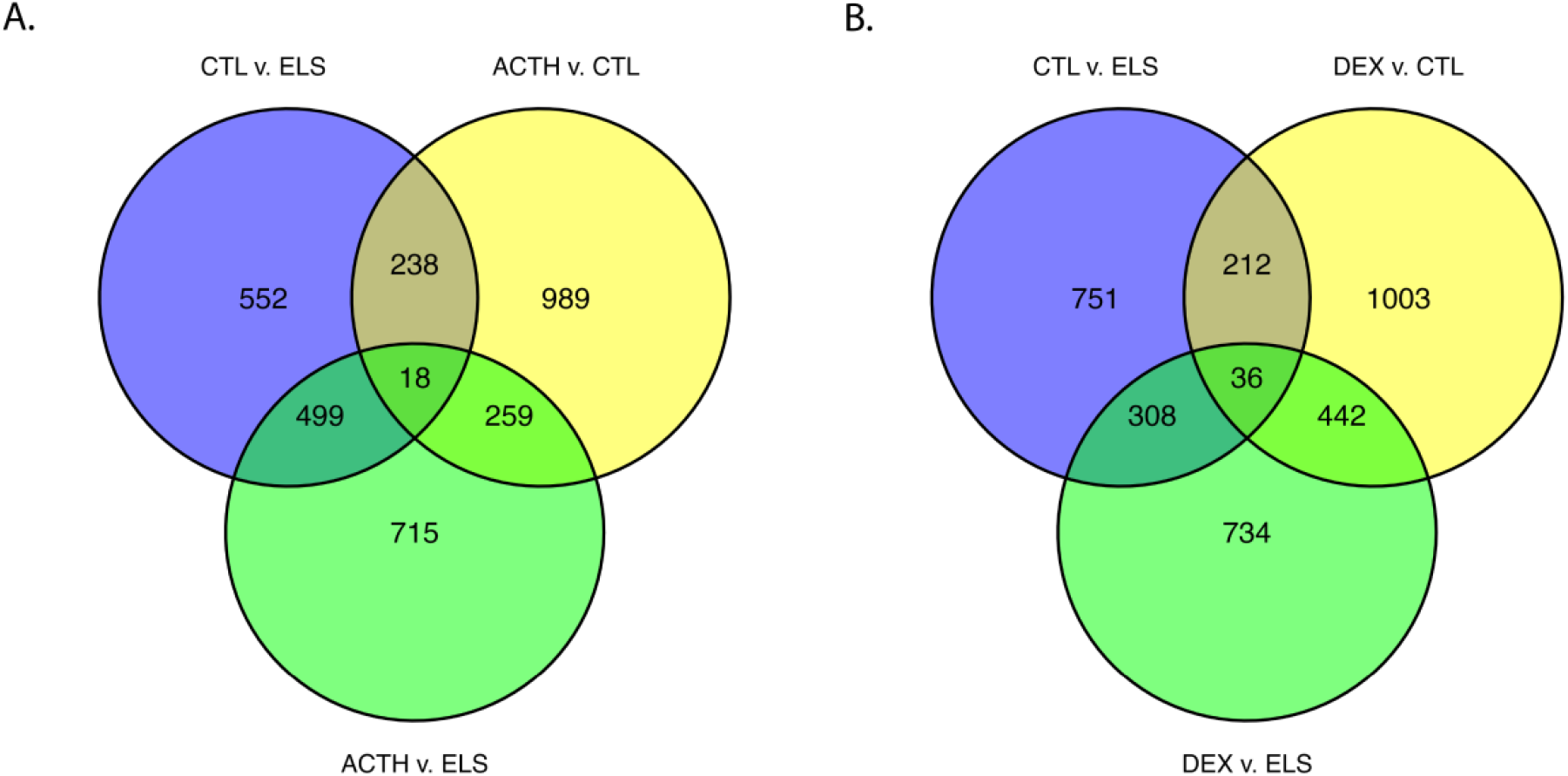
Venn diagrams showing the overlap of DEGs between the different model contrasts. A) The plot shows the distribution of shared and unique DEGs between the ACTH vs. Control (CTL), ACTH vs. vehicle ELS (ELS) and Control vs. vehicle ELS contrasts. B) The plot shows the distribution of shared and unique DEGs between the Dex vs. Control (CTL), Dex vs. vehicle ELS (ELS) and Control vs. vehicle ELS contrasts. There was a significantly greater number of DEGs shared between the CTL vs. ELS and ACTH vs. ELS than shared between the CTL vs. ELS and Dex vs. ELS contrasts (Chi-squared p = 10e-13) indicating ACTH after seizure induction helps normalize some of the genetic signal back to control levels.

We performed pathway analysis to identify the enriched pathways that are differentially expressed in response to each drug to counteract or exacerbate cellular and molecular dysfunction caused by ELS. We performed ontology analysis on the shared set of DEGs of the Control v. ELS, and Dex and ACTH v. ELS contrasts. Although there were 344 shared genes between Control and Dex, only a single pathway was enriched: “regulation of transcription elongation from RNA polymerase II promoter” (GO:0034243) (Fig 2B). To identify the pathways that are dysregulated by ELS but normalized by ACTH, we performed ontology analysis on the shared set of genes between the ACTH vs. vehicle ELS and Control vs. vehicle ELS contrasts. The analysis identified twenty-four enriched terms including several that were brain-specific and tightly related to neuronal and glial dynamics such as “regulation of synaptic plasticity” (GO:0048167) and “supramolecular fiber organization” (GO:0097435) (Fig 2A). Taken together, these terms in the ACTH-treated group broadly relate to cell survival and to homeostatic processes that are targeted by ACTH.

**Figure 2:**
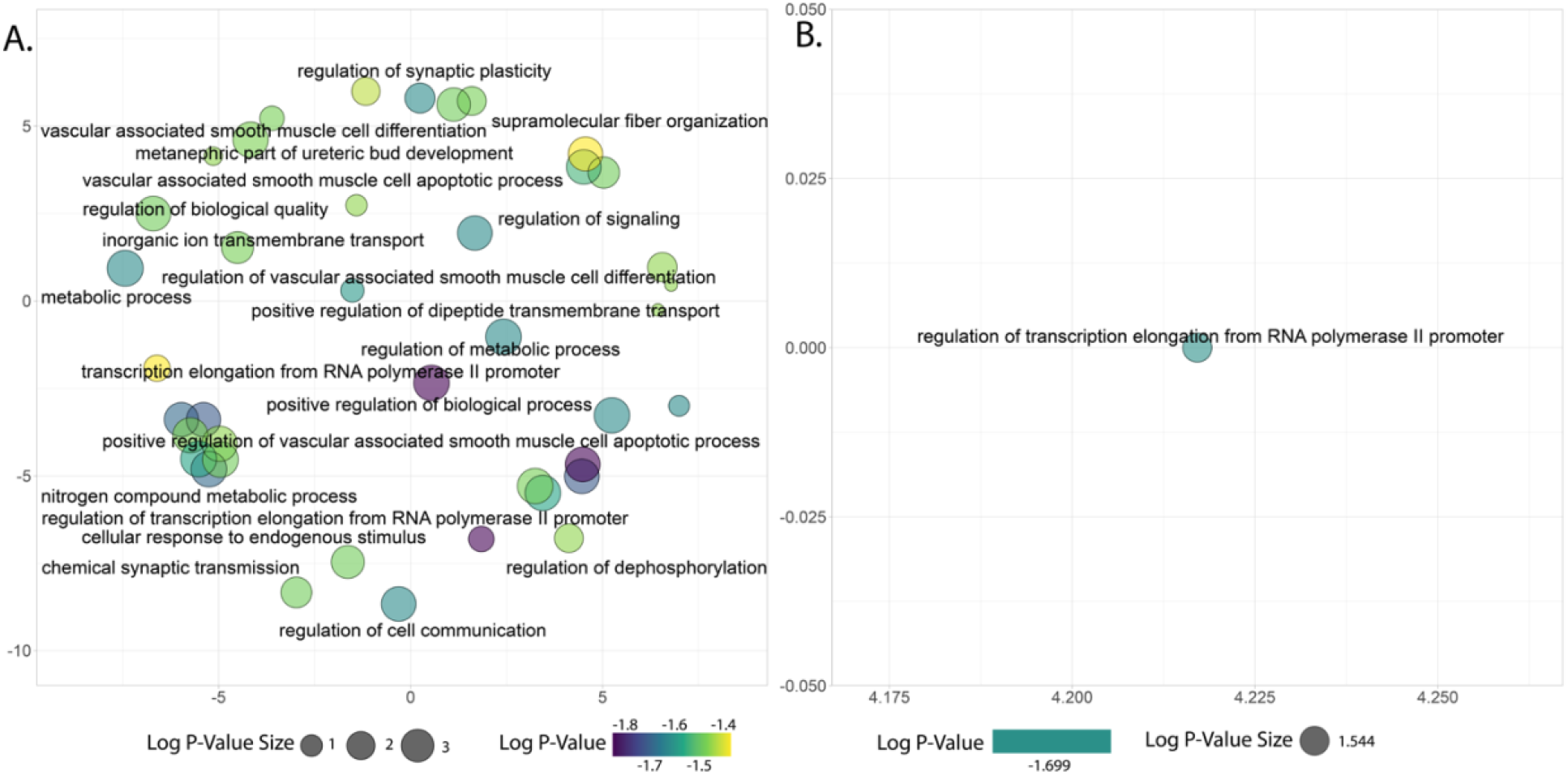
Semantic plots representing the diversity of ontology terms enriched by DEGs. A) Enrichments from the set of DEGs shared between the ACTH vs. vehicle ELS contrast and the Control vs. vehicle ELS contrast. B) The single enriched pathway from the set of DEGs shared between the Dex vs. vehicle ELS contrast and the Control vs. vehicle ELS contrast.

These results together indicate that while Dex in the context of ELS appears to influence the expression of a larger proportion of genes, it does little to shift gene expression back toward the Control signal. ACTH in the context of ELS appears to influence a more specific subset of genes to move back toward the “normal” expression of the non-ELS Control animals.

### ACTH alters expression of genes involved in synaptic regulation post-ELS

We next widened our scope and performed pathway enrichment analysis for the whole set of DEGs from each model contrast. We first examined the shared pathway enrichments of animals treated with ACTH and Dex (vs. ELS) to the enrichments of Control (vs. ELS) animals (Fig. 3A&B). Each contrast has a subset of unique pathways that are solely enriched but they also share a number of functional pathways as well. The enriched pathways shared between ACTH and Control are involved in the proper regulation of ion transport, as well as intracellular and trans-synaptic signaling (Fig 3A). These pathway enrichments may be in response to dysregulated ion transport post-seizure. We examined the set of ion-transport DEGs and within the top 10 there were several genes involved in the regulation of Na^+^ and K^+^ balance (*Aqp8, Fxyd7, Fgf12*) and Ca2+ regulation (*Fgf12, Mchr1, Syt10*). This suggests that in the context of ELS, a treatment with ACTH might regulate ion dynamics at the synapse post-seizure. Human orthologs of genes like *Fgf12* and *Syt10* have been previously associated with different epilepsies. Specifically, *Fgf12* negatively regulates Ca^2+^ channels to regulate membrane depolarization and intracellular calcium dependent pathways. Up-regulation of this gene by ACTH might help attenuate network excitability and calcium neurotoxicity post-seizure.

**Figure 3:**
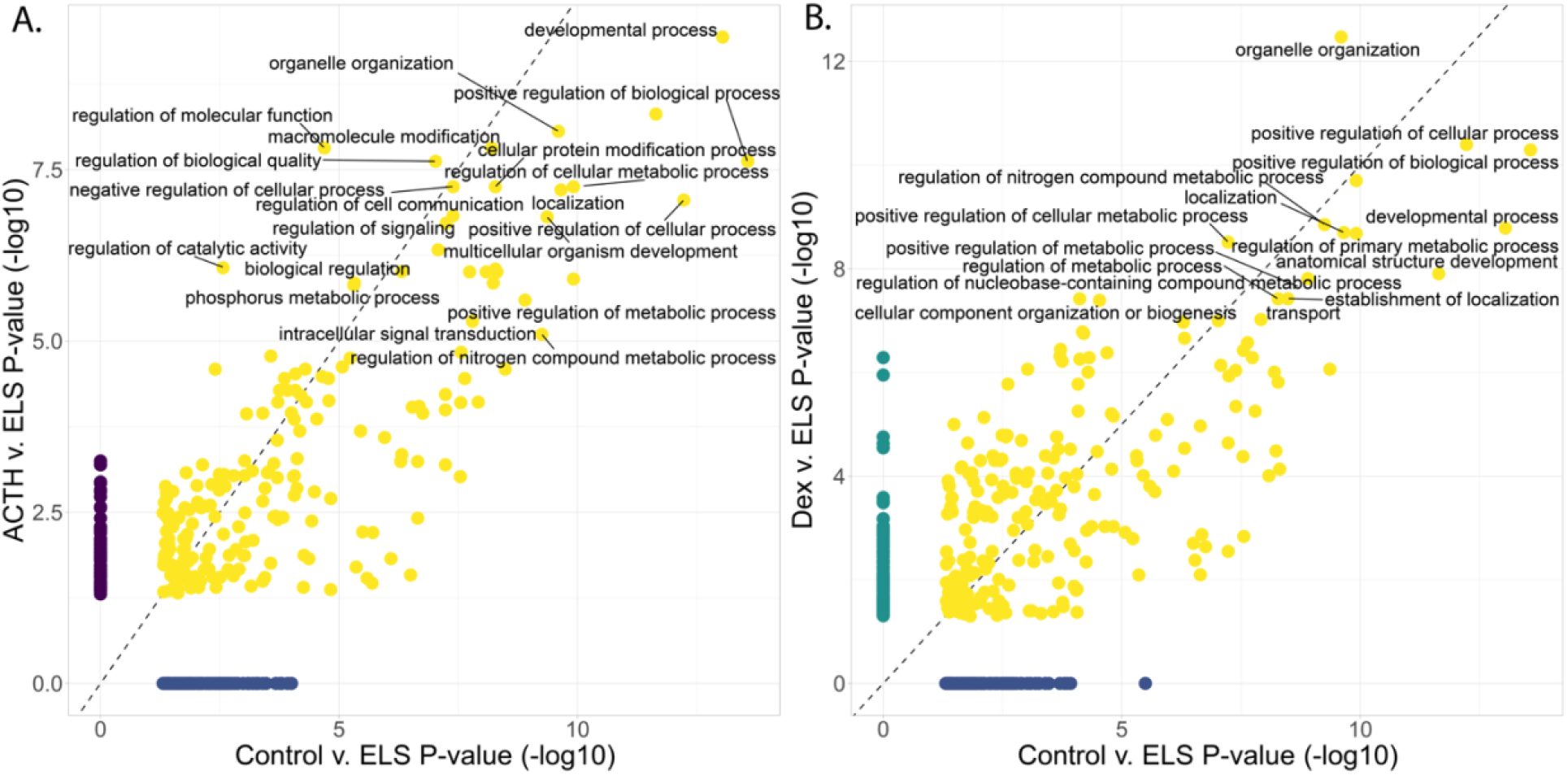
Comparison of enrichments from contrasts compared to ELS. A) Ontology comparison between the ACTH vs. vehicle ELS and Control vs. vehicle ELS contrasts. Several relevant pathways are enriched by ACTH treatment and in the Control group such as “regulation of ion-transport”. This supports the idea that ACTH is influencing expression towards Control patterns in the context of ELS. B) Ontology comparison between the Dex vs. vehicle ELS and Control vs. vehicle ELS contrasts. The shared enrichments here are more general and suggest that Dex is less effective at targeting pathways that could improve cognitive outcomes in ELS individuals.

The picture painted by the shared Dex and Control pathway analysis is much less precise (Fig 3B). While we observed terms like “Response to Stress”, which could indicate the enrichment of genes to respond to the stressful cellular environments post-ELS, the rest of the terms are more general and tend to be involved in regulating metabolic processes. While these enrichments make sense for a steroid like Dex, they seem more scattershot, and not as specifically involved with neuronal or glial recovery post-ELS.

To explore the pathway enrichments more deeply, we next compared the shared and unique pathway enrichments of Dex and ACTH animals (Fig. 4A-C). Figure 4A indicates the enriched pathways shared by the two treatment groups. We observed a wide range of shared pathway enrichments from regulation of metabolic processes to the regulation of synaptic plasticity and trans-synaptic signaling. This indicates that both drugs do work in similar pathways and can both influence seizure-relevant pathways. Taken together with the findings above, support the hypothesis that Dex exerts more systemic effects while ACTH exerts more localized CNS effects.

**Figure 4:**
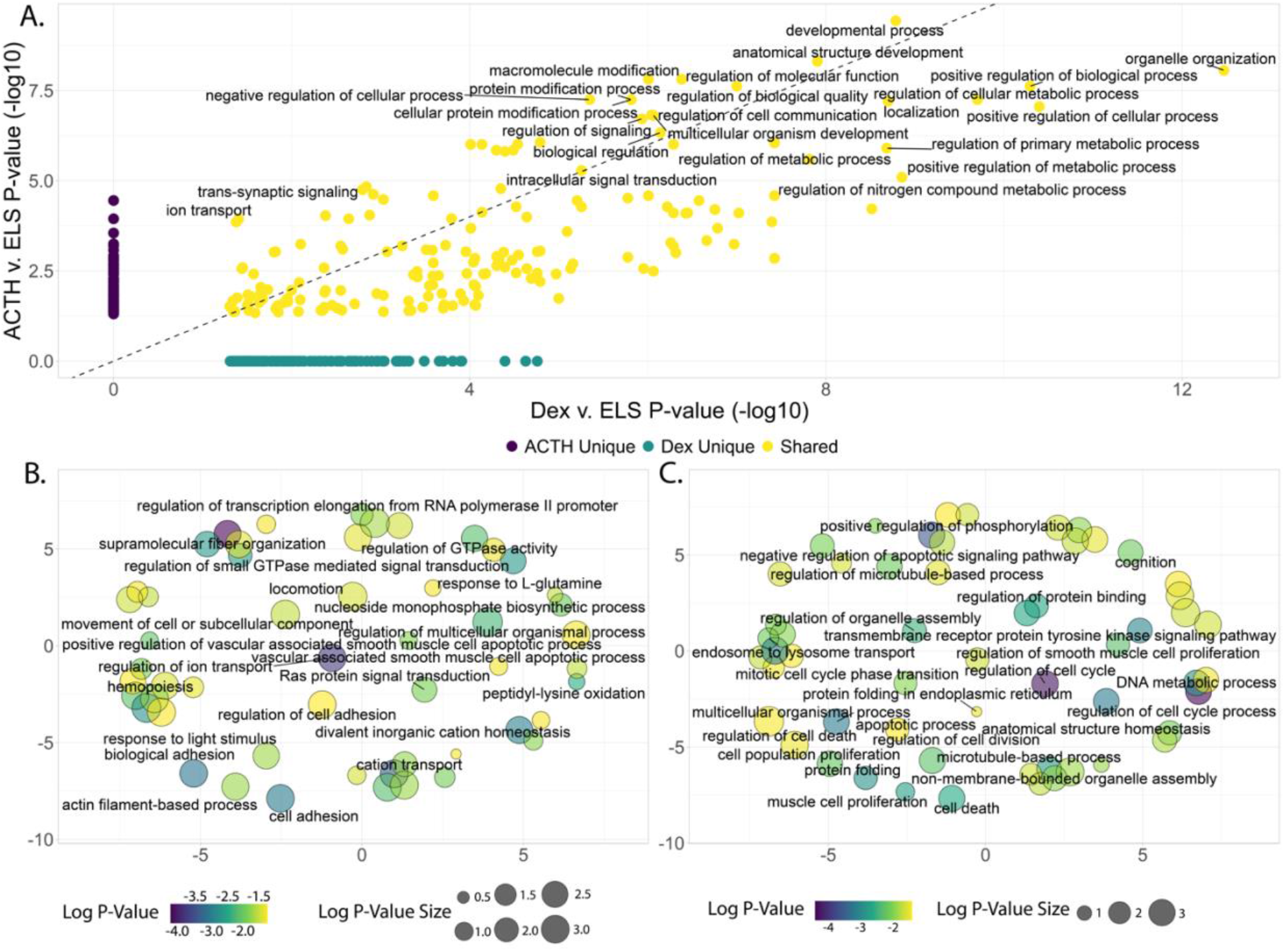
Direct comparison of ACTH vs. vehicle ELS and Dex vs. vehicle ELS contrast enrichments. A) The shared set of enrichments for ACTH and DEX compared to ELS are colored in yellow. Several of the enrichments are involved in seizure-related pathways indicating that Dex is working in some ELS-relevant pathways. B) Unique enrichments (colored in purple in A) for ACTH vs. vehicle ELS. Pathways like regulation of ion transport and supramolecular fiber organization have been previously noted to play important roles in post-ELS molecular and cellular response. C) Unique enrichments (colored in green in A) for Dex vs. vehicle ELS. These terms are more general, showing that while Dex is working in some relevant pathways, it is less precise in its regulation of seizure-relevant pathways.

We next examined the unique pathway enrichments for each treatment group. The ACTH group had fewer unique enriched pathways (n = 128) (Fig 4B) than the Dex group (n = 170) (Fig 4C). However, while the Dex treated animals still had unique enrichments involved in synaptic signaling, Dex also influenced many other pathways seemingly unrelated to the ELS insult. ACTH on the other hand, with its fewer unique pathway enrichments, nevertheless has a more precise set of functional pathways involved in post-ELS amelioration processes with enrichments in pathways like “glial cell proliferation”, “trans-synaptic signaling”, and “supramolecular fiber organization”, among several others.

The pathway enrichments further underscore that while Dex is affecting a broad range of gene expression and influencing numerous functional pathways, in the case of ELS it is less specific, whereas ACTH drives more specific changes in response to the ELS insult.

## Discussion

Complex seizure disorders are caused by a multitude of genetic variations and environmental insults. The identification of differentially regulated genes and pathways will help us better understand the underlying dysfunctional mechanisms of seizure disorders and may reveal the best paths forward for effectively treating their negative outcomes. Our overall goal is to elucidate the genetics contributing to negative outcomes and eventual improvement of those outcomes via ACTH or Dex. The methodological setup, however, made it impossible to directly link behavioral changes to the genetic ones we observed. The main focus of the project was determining differential expression after early intervention with ACTH and Dex which required rapid extraction of the brain tissue for sequencing, prohibiting any behavioral tests. We have instead relied upon our previous behavioral observations in rats treated with ACTH and Dex as described in the introduction to inform the significance of our genetic findings (Massey et al., 2016).

We hypothesized that ACTH would ameliorate the negative impact of ELS by returning gene expression to a more normal pattern. We observed that ACTH treatment resulted in a gene expression pattern with fewer DEGs compared to Controls than Dex. Additionally, when compared to vehicle-treated animals that underwent ELS, animals treated with ACTH and Control animals shared a much larger proportion of DEGs than Dex and Control animals. This, despite Dex’s larger proportion of DEGs compared to ELS, suggests that ACTH treatment normalizes gene expression in a more targeted way. Furthermore, while ACTH possessed a smaller proportion of unique significantly enriched pathways compared to Dex, those that were enriched were involved in pathways that are likely to be involved in post-seizure recovery such as peptidyl-amino acid modification and regulation of DNA metabolic processes. It appears that while Dex has broad influence over a diverse range of functional pathways in the brain, ACTH is precisely modifying pathways involved in CNS recovery and returning the gene expression landscape to normal.

Though the gene expression signature for ACTH is closer to that of Controls, it does not perfectly reverse the effects of ELS. However, the genes ACTH does influence back towards Control patterns of expression seem to be important for the regulation of crucial neuro-regulatory pathways such as ion balance, neuronal signaling, and glial cell proliferation. Altering the expression of key genes in these pathways could help reduce the hyperexcitability of the system and ameliorate cognitive deficits that result from the molecular and genetic dysfunction of seizure disorders.

The “regulation of ion transport” (GO:0043269) GO term is of particular interest as ion dysregulation is a key feature of excitatory/inhibitory balance in seizure disorders (Fritschy, 2008). Repetitive firing of neurons increases the extracellular concentrations of K^+^. This leads to altered extracellular ion concentrations which can make the surrounding neurons more hyperexcitable. The specific genes enriched by ACTH in the context of ELS seem to oppose this hyperexcitability by controlling ionic concentrations of K^+^, Na^+^ and Ca^2+^.

While it has yet to be studied in the context of seizure disorders, Synaptotagmin10 (*Syt10*) has recently been observed to be a key mediator of excitotoxicity in a model of neurodegeneration (Woitecki et al., 2016). The group observed that *Syt10* is required for the protection of neurons in the hippocampus and seems to be part of the core sets of genes involved in neuroprotection against brain insults.

The gene *Fgf12* which is involved in the negative regulation of cation channel activity (Wildburger et al., 2015). Mutations in the human ortholog, *FHF1*, have led to rare epileptic encephalopathies (Al-Mehmadi et al., 2016), possibly by hampering the ability of the protein to associate with the C terminal tails of sodium channels. Upregulation of this gene by ACTH post-ELS could help temper hyperexcitable neurons by lowering opening probability of sodium channels which would lower chances of cell depolarization. In contrast, the gene *Ptgs2* (*Cox2*) is downregulated by ACTH post-ELS. This gene has been actively studied in the context of seizure disorders and other neurological diseases for the past two decades (Okada et al., 2001; Chen et al., 2002; Almalki et al., 2014). In 2002, Chen and colleagues observed that selective *COX2* inhibitors decreased membrane excitability and decreased back propagation action potential induced Ca2+ influx, a key feature of LTP. Taken together, normalization of these genes’ expression by ACTH may underlie better control of calcium homeostasis and hyperexcitability that is required for normal neural network function and subsequently preventing cognitive dysfunction; as such, this may be part of the molecular mechanisms for improved outcome seen with ACTH treatment (Sher and Sheikh, 1993; Hernan et al., 2014; Altunel et al., 2017b, 2017a).

Another unique term enriched by DEGs in the context of ACTH is “supramolecular fiber organization” (GO:0097435). Post-insult, neuronal and glial cells undergo changes to their actin filaments as glia move to respond to injured tissue and dendritic branches are pruned and rearranged as neurons respond to neighboring damaged cells (Jackson et al., 2012). The fiber organization genes enriched by ACTH in the context of ELS are involved in a myriad of processes including combatting cell death and promoting neuroprotective pathways. The gene *Ilk* is an integrin-linked kinase which negatively regulates apoptosis and has previously been shown to be downregulated post-pilocarpine-induced status epilepticus (Kim et al., 2009). In this study, *Ilk* is upregulated in ACTH compared to ELS. The gene *Jmy* was shown to inhibit neuritogenesis at high levels of expression (Firat-Karalar et al., 2011). An increased expression of *Jmy* may temper changes in neuronal connections post-seizure, by preventing new connections to form between neurons, that may contribute to neuronal hyperexcitability in the network. In contrast to these upregulated genes, *Fat1*, (human ortholog *FAT1*) has been implicated in autism spectrum disorder (ASD) in humans and is downregulated in the context of ACTH (Frei et al., 2021). Less is known about the functional role of this gene at this point in time. It has been identified as a tumor suppressor and is predicted to act upstream of actin filament organization and cell-cell adhesion (Yaoita et al., 2005; Fang et al., 2019). While less is known about the precise functional roles of *Fat1*, increased mobility of actin and lowering of cell-cell adhesion could make it easier for microglia to migrate to damaged areas, to remove damaged neurons. Both *Ilk* and *Jmy* are upregulated in the context of ACTH compared to ELS. Previous work in the literature has reported that higher levels of expression in both of these genes promotes beneficial pathways for neuroprotection and cell-survival.

Proposed mechanisms of action for ACTH generally involve binding of ACTH to melanocortin 2 receptors (MC2Rs) in the adrenal cortex, leading to downstream release of corticosteroids. We have previously shown that ACTH and Dex, a corticosteroid, lead to different outcomes in terms of cognition and in this study to differential gene expression. One hypothesis for the action of ACTH above and beyond those of Dex is the presence of other subtypes of the melanocortin receptors, MC1Rs, MC3Rs, and MC4Rs, in the CNS. MC4Rs in particular have been studied for their role in neuroprotection in other disease states. MC4Rs are found on many cell types in the brain, consistent with the glial- and neuron-specific gene expression pathways. Binding of melanocortins like ACTH centrally to glial cell MC4Rs can have anti-neuroinflammatory effects (Catania, 2008), while binding of melanocortins to neuronal MC4Rs can directly modulate neuronal firing and synaptic plasticity (Shen et al., 2013). We have indication that genes in both of these pathways are differentially expressed in animals treated with ACTH, but not those treated with Dex (Catania, 2008; Shen et al., 2013). Mounting evidence has found that activation of MC4R in particular can decrease cell death in a number of disease models, improve cognitive outcomes in models of Alzheimer’s disease and cerebral ischemia, and can regulate synaptic plasticity in the hippocampus (Aronsson et al., 2007; Spaccapelo et al., 2011; Giuliani et al., 2014; Ma and McLaurin, 2014; Shen et al., 2016). Our results in this study indicate that the genes and pathways ACTH influences help push gene expression in the brain back to a normal state of expression by upregulating genes which support homeostatic balance and downregulating those which might contribute to excitotoxic cell damage post-ELS; in support of the more targeted and brain-specific differential gene expression, we suggest that ACTH may exert its differential actions through pathways that are dependent on melanocortin receptors in the brain.

## Acknowledgements

We would like to acknowledge the Vermont Integrated Genomics Core (VIGC) for their help in processing the sequencing data and the Vermont Advaced Computing Core (VACC) for providing access to their high-performance computing cluster.

## Figure legends

Table 1-1: Differentially expressed genes for the Control vs. vehicle ELS contrast.

Table 1-2: Differentially expressed genes for the ACTH vs. Control contrast.

Table 1-3: Differentially expressed genes for the ACTH vs. vehicle ELS contrast.

Table 1-4: Differentially expressed genes for the Dex vs. Control contrast.

Table 1-5: Differentially expressed genes for the Dex vs. vehicle ELS contrast.

Table 2-1: Significantly enriched GO terms shared between the ACTH vs. vehicle ELS and Control vs. vehicle ELS contrasts.

Table 2-2: Significantly enriched GO terms shared between the Dex vs. vehicle ELS and Control vs. vehicle ELS contrasts.

Table 3-1: Significantly enriched Go terms for the Control vs. vehicle ELS contrast.

Table 4-1: Significantly enriched GO terms for the ACTH vs. vehicle ELS contrast.

Table 4-2: Significantly enriched GO terms for the Dex vs. vehicle ELS contrast

## Abbreviations

(ELS): Early-life seizures
(ACTH): adrenocorticotropic hormone
(Dex): dexamethasone
(DEG): differential gene expression

## References

Almalki A, Alston CL, Parker A, Simonic I, Mehta SG, He L, Reza M, Oliveira JMA, Lightowlers RN, McFarland R, Taylor RW, Chrzanowska-Lightowlers ZMA (2014) Mutation of the human mitochondrial phenylalanine-tRNA synthetase causes infantile-onset epilepsy and cytochrome c oxidase deficiency. Biochimica Et Biophysica Acta Bba - Mol Basis Dis 1842:56–64.

Al-Mehmadi S, Splitt M, group* FDS, Ramesh V, DeBrosse S, Dessoffy K, Xia F, Yang Y, Rosenfeld JA, Cossette P, Michaud JL, Hamdan FF, Campeau PM, Minassian BA, group‡ FCenS, group DS, Barrett J, Hurles M (2016) FHF1 (FGF12) epileptic encephalopathy. Neurology Genetics 2:NA;

Altunel A, Altunel EÖ, Sever A (2017a) Response to adrenocorticotropic in attention deficit hyperactivity disorder-like symptoms in electrical status epilepticus in sleep syndrome is related to electroencephalographic improvement: A retrospective study. Epilepsy Behav 74:161–166.

Altunel A, Sever A, Altunel EÖ (2017b) ACTH has beneficial effects on stuttering in ADHD and ASD patients with ESES: A retrospective study. Brain Dev 39:130–137.

Anders S, Pyl PT, Huber W (2015) HTSeq—a Python framework to work with high-throughput sequencing data. Bioinformatics 31:166–169.

Aronsson ÅF, Spulber S, Oprica M, Winblad B, Post C, Schultzberg M (2007) α-MSH Rescues Neurons from Excitotoxic Cell Death. J Mol Neurosci 33:239.

Bentley DR et al. (2008) Accurate whole human genome sequencing using reversible terminator chemistry. Nature 456:53–59.

Catania A (2008) Neuroprotective actions of melanocortins: a therapeutic opportunity. Trends Neurosci 31:353–360.

Chen C, Magee JC, Bazan NG (2002) Cyclooxygenase-2 Regulates Prostaglandin E2 Signaling in Hippocampal Long-Term Synaptic Plasticity. J Neurophysiol 87:2851–2857.

Dobin A, Davis CA, Schlesinger F, Drenkow J, Zaleski C, Jha S, Batut P, Chaisson M, Gingeras TR (2013) STAR: ultrafast universal RNA-seq aligner. Bioinformatics 29:15–21.

Duffy BA, Chun KP, Ma D, Lythgoe MF, Scott RC (2014) Dexamethasone exacerbates cerebral edema and brain injury following lithium-pilocarpine induced status epilepticus☆. Neurobiol Dis 63:229–236.

Dunn DW, Kronenberger WG (2005) Childhood Epilepsy, Attention Problems, and ADHD: Review and Practical Considerations. Semin Pediatr Neurol 12:222–228.

Fang W, Ma Y, Yin JC, Hong S, Zhou H, Wang A, Wang F, Bao H, Wu X, Yang Y, Huang Y, Zhao H, Shao YW, Zhang L (2019) Comprehensive Genomic Profiling Identifies Novel Genetic Predictors of Response to Anti–PD-(L)1 Therapies in Non–Small Cell Lung Cancer. Clin Cancer Res 25:5015–5026.

Firat-Karalar EN, Hsiue PP, Welch MD (2011) The actin nucleation factor JMY is a negative regulator of neuritogenesis. Mol Biol Cell 22:4563–4574.

Frei JA, Brandenburg CJ, Nestor JE, Hodzic DM, Plachez C, McNeill H, Dykxhoorn DM, Nestor MW, Blatt GJ, Lin Y-C (2021) Postnatal expression profiles of atypical cadherin FAT1 suggest its role in autism. Biol Open 10:bio056457.

Fritschy J-M (2008) Epilepsy, E/I Balance and GABAA Receptor Plasticity. Front Mol Neurosci 1:5.

Ghosh C, Myers R, O’Connor C, Williams S, Liu X, Hossain M, Nemeth M, Najm IM (2022) Cortical Dysplasia in Rats Provokes Neurovascular Alterations, GLUT1 Dysfunction, and Metabolic Disturbances That Are Sustained Post-Seizure Induction. Mol Neurobiol:1–18.

Giuliani D, Bitto A, Galantucci M, Zaffe D, Ottani A, Irrera N, Neri L, Cavallini GM, Altavilla D, Botticelli AR, Squadrito F, Guarini S (2014) Melanocortins protect against progression of Alzheimer’s disease in triple-transgenic mice by targeting multiple pathophysiological pathways. Neurobiol Aging 35:537–547.

Hernan AE, Alexander A, Lenck-Santini P-P, Scott RC, Holmes GL (2014) Attention Deficit Associated with Early Life Interictal Spikes in a Rat Model Is Improved with ACTH. Plos One 9:e89812.

Jackson J, Chugh D, Nilsson P, Wood J, Carlström K, Lindvall O, Ekdahl CT (2012) Altered Synaptic Properties During Integration of Adult-Born Hippocampal Neurons Following a Seizure Insult. Plos One 7:e35557.

Kanner AM (2005) Depression in Epilepsy: A Neurobiologic Perspective. Epilepsy Curr 5:21–27.

Kim GW, Kim H-J, Cho K-J, Kim H-W, Cho Y-J, Lee BI (2009) The role of MMP-9 in integrin-mediated hippocampal cell death after pilocarpine-induced status epilepticus. Neurobiol Dis 36:169–180.

Kolberg L, Raudvere U, Kuzmin I, Vilo J, Peterson H (2020) gprofiler2 -- an R package for gene list functional enrichment analysis and namespace conversion toolset g:Profiler. F1000research 9:ELIXIR–709.

Love MI, Huber W, Anders S (2014) Moderated estimation of fold change and dispersion for RNA-seq data with DESeq2. Genome Biol 15:550.

Ma K, McLaurin J (2014) α-Melanocyte Stimulating Hormone Prevents GABAergic Neuronal Loss and Improves Cognitive Function in Alzheimer’s Disease. J Neurosci 34:6736–6745.

Martin M (2011) Cutadapt Removes Adapter Sequences From High-Throughput Sequencing Reads. EMBnet.journal 17:10–12.

Massey AT, Lerner DK, Holmes GL, Scott RC, Hernan AE (2016) ACTH Prevents Deficits in Fear Extinction Associated with Early Life Seizures. Front Neurol 7:65.

Matson JL, Neal D, Hess JA, Mahan S, Fodstad JC (2010) The effect of seizure disorder on symptom presentation in atypically developing children and children with autism spectrum disorders based on the BDI-2. Dev Neurorehabil 13:310–314.

Nariai H, Duberstein S, Shinnar S (2018) Treatment of Epileptic Encephalopathies: Current State of the Art. J Child Neurol 33:41–54.

Okada K, Yuhi T, Tsuji S, Yamashita U (2001) Cyclooxygenase-2 expression in the hippocampus of genetically epilepsy susceptible El mice was increased after seizure. Brain Res 894:332–335.

Rantanen K, Nieminen P, Eriksson K (2010) Neurocognitive functioning of preschool children with uncomplicated epilepsy. J Neuropsychol 4:71–87.

Shen Y, Fu W-Y, Cheng EYL, Fu AKY, Ip NY (2013) Melanocortin-4 Receptor Regulates Hippocampal Synaptic Plasticity through a Protein Kinase A-Dependent Mechanism. J Neurosci 33:464–472.

Shen Y, Tian M, Zheng Y, Gong F, Fu AKY, Ip NY (2016) Stimulation of the Hippocampal POMC/MC4R Circuit Alleviates Synaptic Plasticity Impairment in an Alzheimer’s Disease Model. Cell Reports 17:1819–1831.

Sher PK, Sheikh MR (1993) Therapeutic efficacy of ACTH in symptomatic infantile spasms with hypsarrhythmia. Pediatr Neurol 9:451–456.

Spaccapelo L, Bitto A, Galantucci M, Ottani A, Irrera N, Minutoli L, Altavilla D, Novellino E, Grieco P, Zaffe D, Squadrito F, Giuliani D, Guarini S (2011) Melanocortin MC4 receptor agonists counteract late inflammatory and apoptotic responses and improve neuronal functionality after cerebral ischemia. Eur J Pharmacol 670:479–486.

Supek F, Bošnjak M, Škunca N, Šmuc T (2011) REVIGO Summarizes and Visualizes Long Lists of Gene Ontology Terms. Plos One 6:e21800.

Wildburger NC, Ali SR, Hsu W-CJ, Shavkunov AS, Nenov MN, Lichti CF, LeDuc RD, Mostovenko E, Panova-Elektronova NI, Emmett MR, Nilsson CL, Laezza F (2015) Quantitative Proteomics Reveals Protein-Protein Interactions with Fibroblast Growth Factor 12 as a Component of the Voltage-Gated Sodium Channel 1.2 (Nav1.2) Macromolecular Complex in Mammalian Brain*. Mol Cell Proteomics 14:1288–1300.

Woitecki AMH, Müller JA, Loo KMJ van, Sowade RF, Becker AJ, Schoch S (2016) Identification of Synaptotagmin 10 as Effector of NPAS4-Mediated Protection from Excitotoxic Neurodegeneration. J Neurosci 36:2561–2570.

Yaoita E, Kurihara H, Yoshida Y, Inoue T, Matsuki A, Sakai T, Yamamoto T (2005) Role of Fat1 in cell-cell contact formation of podocytes in puromycin aminonucleoside nephrosis and neonatal kidney. Kidney Int 68:542–551.

Yates AD et al. (2019) Ensembl 2020. Nucleic Acids Res 48:D682–D688.

